# Preserved spontaneous mentalizing amid reduced intersubject variability in autism during a movie narrative

**DOI:** 10.1101/2024.03.08.583911

**Authors:** Margot Mangnus, Saskia B. J. Koch, Kexin Cai, Miriam Greidanus Romaneli, Peter Hagoort, Jana Bašnáková, Arjen Stolk

**Affiliations:** Donders Institute for Brain, Cognition, and Behaviour, Radboud University, Nijmegen, the Netherlands; Psychological and Brain Sciences, Dartmouth College, Hanover, NH, USA; Max Planck Institute for Psycholinguistics, Nijmegen, the Netherlands

**Author notes:** These authors contributed equally (Jana Bašnáková, Arjen Stolk). Corresponding author Margot Mangnus Donders Centre for Cognitive Neuroimaging P.O. Box 9101, NL-6500 HB, Nijmegen, the Netherlands.

**Keywords:** mentalizing, intersubject correlation, social neuroscience, naturalistic stimuli

## Abstract

Individuals with autism may perform well in structured Theory of Mind (ToM) tasks that assess their ability to infer mental states, yet encounter challenges in everyday social interactions. Using fMRI and pupillometry, we investigated whether this discrepancy stems from reduced spontaneous mentalizing or broader difficulties in unstructured environments. Autistic adults and neurotypical controls viewed a nonverbal animated movie featuring events known to evoke mental state inferences. Both groups exhibited similar ToM network activations and pupil responses to these events, alongside comparable verbal accounts of the mental states portrayed. However, intersubject correlation analysis revealed a significant reduction in response variability among autistic participants throughout the movie’s complex visual narrative, affecting brain regions beyond the ToM network. We suggest that the primary challenge for individuals with autism may lie in the idiosyncratic exploration of narratives in unstructured settings, regardless of whether mental state inferences are involved.

## Introduction

Autism Spectrum Condition (ASC) is a neurodevelopmental disorder marked by difficulties in social communication and interaction across multiple contexts (American Psychiatric Association, 2013). These difficulties are frequently associated with Theory of Mind (ToM) – the ability to attribute mental states to oneself and others, also known as mentalizing (Premack & Woodruff, 1978; Wimmer & Perner, 1983). While initial findings suggested diminished ToM abilities in autism (Baron-Cohen et al., 1985; Happé, 1994), subsequent studies have painted a more complex picture, owing to a variety of factors.

First, the reliability of established ToM assessments, such as the False Belief Test, Strange Stories, and the Reading the Mind in the Eye Test, has been increasingly questioned. These tests exhibit inconsistent effect sizes when compared to earlier, smaller-scale studies and show limited correlation with one another despite their aim of measuring the same ToM construct (Gernsbacher & Yergeau, 2019; Higgins et al., 2024; Schaafsma et al., 2015). Second, studies often fail to match autistic individuals with neurotypical controls based on language abilities in explicit tasks, i.e., requiring verbal responses or comprehension. This lack in language matching is a critical factor considering the generally lower performance in verbal learning and memory among the autism population (Velikonja et al., 2019). Lastly, recent research underscores the remarkable proficiency of autistic individuals in ToM tasks, especially in situations requiring strategic mental state reasoning (Bowler, 1992; Pantelis & Kennedy, 2017), such as deception (van Tiel et al., 2021).

Faced with these empirical challenges, researchers have turned to more implicit ToM measures. This includes the analysis of anticipatory eye movements concerning an actor’s false beliefs (Senju et al., 2009), and measuring neural activity in the Animated Triangles task, where participants attribute mental states to moving shapes (Abell et al., 2000). Recent findings indicate that while autistic individuals do show anticipatory gaze responses, these responses tend to be generally slower, regardless of whether the character’s beliefs were true or false (Glenwright et al., 2021; Schuwerk et al., 2016). In the Animated Triangles task, both autistic and neurotypical participants show comparable brain activation in key ToM regions, such as the medial prefrontal cortex (mPFC), temporoparietal junction (TPJ), and precuneus (Moessnang et al., 2020). However, individuals with autism typically underperform in both the mentalizing and non-mentalizing conditions of this task, which, combined with their generally slower anticipatory gaze responses, points to broader difficulties in implicit task settings (Wilson, 2021). The extent to which autistic individuals engage in spontaneous mentalizing, particularly in unstructured settings lacking an explicitly defined task, as is common in everyday social interactions, remains largely unknown.

In this study combining fMRI and pupillometry, we sought to bridge this knowledge gap by investigating how individuals with and without autism respond to mental state events embedded in a dynamic movie narrative. Additionally, our aim was to provide insights into how these individuals process stimuli in settings that are less structured and more akin to everyday social interactions. Everyday interactions require the continuous exploration of various stimuli, each imbued with meaning derived from the broader narrative of the interaction (Goffman, 1974; Johnson et al., 2023; Stolk et al., 2022). This is evident in how seemingly inconspicuous behaviors like a cough, a nod, or even silence can carry profound meanings in particular contexts (Kendon, 1994). However, identifying how this narrative processing occurs in unstructured settings is challenging, as it likely unfolds differently over time among individuals exposed to the same stimuli. We reasoned that it could manifest as intersubject variability in responses to movie stimuli, especially among viewers inclined towards exploratory analysis (Chang et al., 2021; Finn et al., 2018; Owen & Manning, 2023; Zhang et al., 2022). We investigated this possibility using intersubject correlation analysis (Hasson et al., 2004), applying it to two complementary methods for assessing cognitive processing; brain imaging and pupil size measurements (Beatty, 1982).

To probe spontaneous mentalizing and narrative processing, we recorded participants’ brain and pupil responses as they viewed the nonverbal animated movie ‘Partly Cloudy’ (Jacoby et al., 2016). Although movies cannot fully replicate the complexities of real-world social interactions (Wheatley et al., 2019), they offer an effective platform for immersing participants in a consistent and evolving narrative under uniform stimulus conditions. To assess differences in narrative processing between autistic and neurotypical participants at various points during the movie, we augmented our intersubject correlation analysis with a dynamic sliding window technique and an adaptive clustering algorithm (Maris & Oostenveld, 2007). The movie ‘Partly Cloudy’ was chosen for its proven ability to evoke neural activations within the ToM network through distinct mental state events (Jacoby et al., 2016; Richardson et al., 2018), making it suitable for evaluating spontaneous mentalizing. After viewing, we analyzed participants’ verbal descriptions of the movie, focusing on their use of language related to mental states. This experimental approach enabled us to simultaneously probe spontaneous mentalizing and narrative processing, providing insights into the interplay of these cognitive functions in individuals with autism.

## Results

### Experimental approach

Fifty-two individuals diagnosed with Autism Spectrum Condition (ASC) and 52 neurotypical (NT) adults participated in our study, matched for gender, age, verbal and nonverbal IQ (Table 1). The ASC participants were all high-functioning, indicative of their relatively lower support needs in daily life. During the experiment, participants underwent simultaneous fMRI scanning and eye-tracking while watching ‘Partly Cloudy’, a Pixar short film previously established as an effective Theory of Mind (ToM) brain network localizer in neurotypical individuals (Jacoby et al., 2016; Paunov et al., 2019). The film’s narrative, centered on the friendship between a stork and a cloud, includes various social interactions. Our analysis focused on events evoking mental state inferences, such as moments where the cloud character contemplates the stork’s actions, and physical state inferences, like the stork experiencing pain (Fig. 1). Control scenes, lacking character interactions, were also included in the analysis. To assess narrative processing during the movie, we employed dynamic intersubject correlation (ISC) analysis. This data-driven approach differs from event-related analyses in that it does not presuppose the relevance of specific movie segments. Additionally, it measures the variability in responses among participants, rather than the overall intensity, to gauge their idiosyncratic engagement with the movie’s narrative.

**Figure 1.**
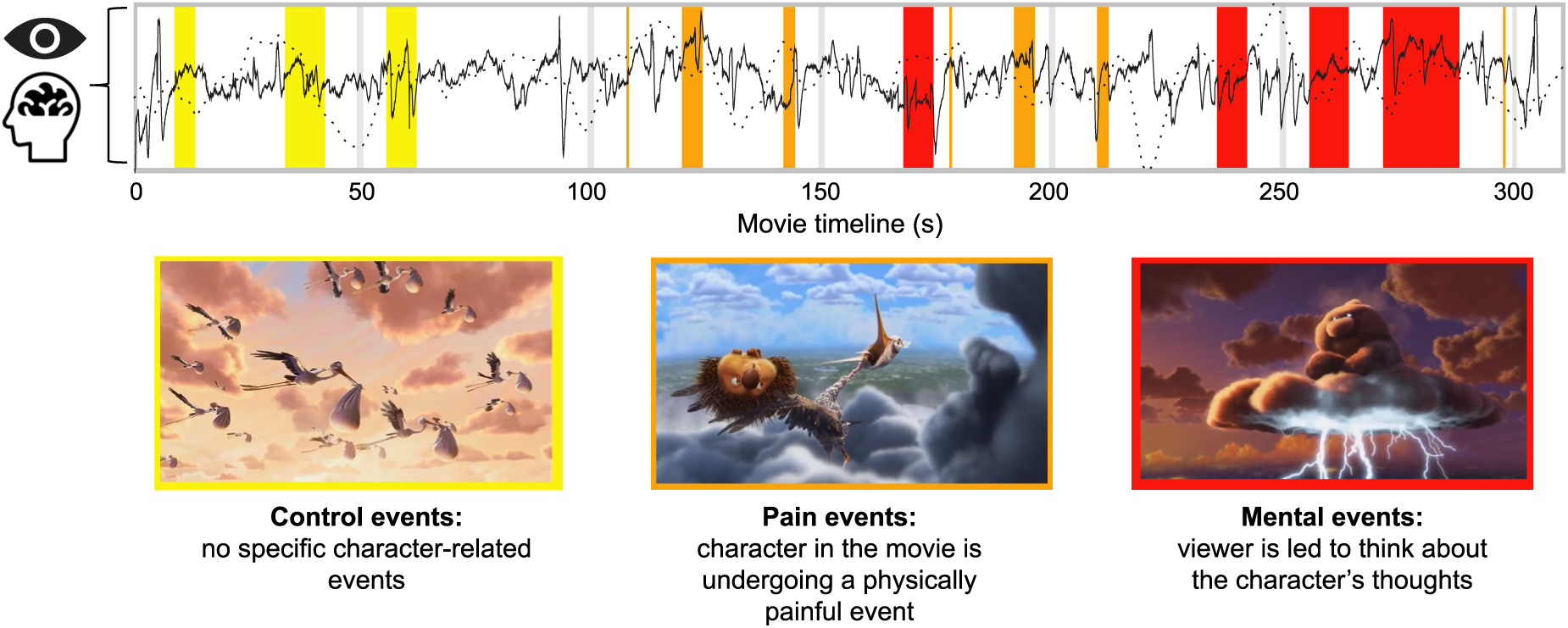
Autistic and neurotypical participants viewed a six-minute animated movie portraying the friendship between a stork and a cloud, while their pupil and brain responses were recorded. Originally employed by Jacoby et al. (2016) as a Theory of Mind localizer, the movie includes three types of events: Mental, Pain and Control, each marked on the movie timeline. Mental and Pain events were expected to prompt inferences about characters’ mental and physical states, whereas Control events featured no characters in the foreground.

**Table 1.** Demographic data.

|  | *ASC Group* | *NT Group* | *Group Difference* |
| --- | --- | --- | --- |
| N (male:female) | 52 (23:29) | 52 (20:32) | $\chi^2 (2, N = 104) = 1.73, p = .19$ |
| Age (years) | 27.7 (6.3) | 26.0 (5.0) | $t (102) = 1.74, p = .08$ |
| Verbal IQ (WAIS-III) | 126 (16) | 124 (16) | $t (102) = 0.51, p = .61$ |
| Nonverbal IQ (RPM) | 103 (9) | 104 (11) | $t (102) = -0.46, p = .64$ |
| Autism Quotient | 31 (9) | 12 (6) | $t (102) = 12.2, p < .001$ |

Values are given as Mean (Standard Deviation). ASC, Autism Spectrum Condition; NT, Neurotypical

### Post-viewing movie descriptions

After the movie, participants were asked to recount the plot and characters’ actions and thoughts in their own words (Fig. 2a). Their verbal descriptions were then evaluated by two independent raters who classified and tallied the words, differentiating between mental state words and other types of content words. The analysis showed no statistically significant difference in the frequency of mental state word usage between autistic and neurotypical participants (M_ASC_ = 0.063, M_NT_ = 0.054, *t*(99) = 0.84, *p* = .40; Fig. 2b), supported by a Bayes Factor in favor of the null hypothesis (BF_Null_ = 3.49). This finding indicates a comparable level of engagement with the mental state events depicted in the movie across both autistic and neurotypical groups.

**Figure 2.**
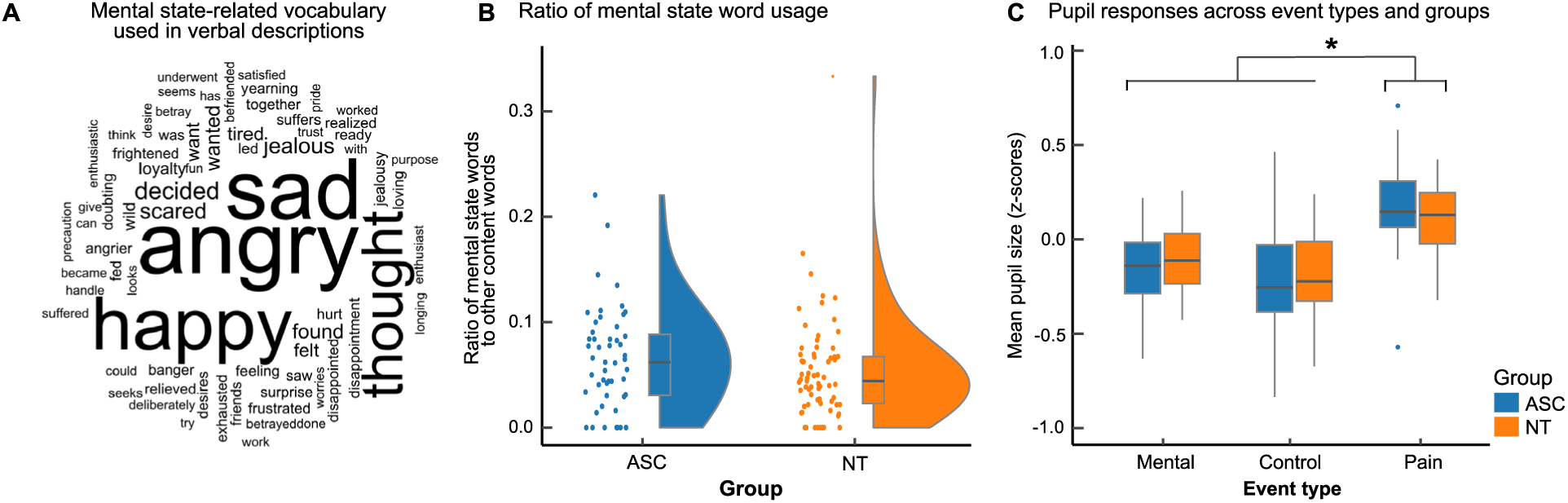
Comparable mental state descriptions and pupil responses across groups. (**a**) Word Cloud depicting the frequency of mental state-related vocabulary used by participants in their post-viewing movie descriptions. The size of each word corresponds to its frequency of use. (**b**) Bar graphs displaying the ratio of mental state words to other content words in these descriptions, showing no significant differences in verbal expressions between the two groups. (**c**) Pupil responses across all event conditions, revealing similar response patterns in both autistic and neurotypical groups. Asterisks denote significant differences between event conditions (all *p* < .001), with no significant differences observed between groups. Outliers are represented by dots, while whiskers display a 1.5 inter-quartile range.

### Event-related pupil responses

Participants’ pupil sizes, considered an index of cognitive processing effort (Beatty, 1982), were continuously tracked throughout the viewing of the movie to assess their responses to mental state events. Analysis of this pupillometric data revealed distinct responses patterns associated with the three main event types (*F*(2,196) = 73.0, *p* < .001, BF_Null_ = 0.00; Fig. 2c). Specifically, *Pain* events induced the largest pupil dilation (M = 0.22, *z*-score), followed by smaller dilations during *Mental* events (M = -0.12) and *Control* events (M = -0.20). Comparing pupil responses between autistic and neurotypical participants revealed no significant differences (*F*(1,98) = 0.03, *p* = .86, BF_Null_ = 7.8). Furthermore, the analysis did not show any significant interaction effects between the groups and event types (*F*(2,196) = 2.1, *p* = .12, BF_Null_ = 2.5), indicating that both autistic and neurotypical individuals exhibited similar pupillary response patterns to the different types of events in the movie.

### Event-related brain responses

Next, we conducted a whole-brain analysis using fMRI to examine neural activation patterns during *Mental* events in comparison to *Control* events. As depicted in Fig. 3a, this analysis revealed robust activation in brain regions associated with the ToM network (Schurz et al., 2014). This included activation in the right and left temporoparietal junction (rTPJ: xyz_MNI_ = [48, -62, 32], t = 16.85, *p_FWE_* < 0.001; lTPJ: [-46, -62, 32], t = 16.92, *p_FWE_* < 0.001), the precuneus ([6, -64, 40], t = 20.64, *p_FWE_* < 0.001), medial prefrontal cortex (mPFC: [-6, 52, 38], t = 7.33, *p_FWE_* < 0.001), and left middle temporal gyrus ([-52, 2, -26], t = 11.51, *p_FWE_* < 0.001). These regions showed increased activation during *Mental* state events compared to *Control* and *Pain* events, emphasizing their role in mental state processing. However, akin to the pupil response findings, there were no significant differences in neural activation patterns between autistic and neurotypical participants, as indicated in both whole-brain and region of interest (ROI) analyses (Fig. 3b). Bayesian statistical approaches corroborated these results, showing a preference for the null hypothesis over models that differentiate between the participant groups. This lack of significant differences was consistently observed across all three ROIs for both *Mental* > *Pain* contrasts (rTPJ: BF_Null_ = 4.83, precuneus: BF_Null_ = 1.73, mPFC: BF_Null_ = 4.01) and *Mental > Control* contrasts (rTPJ: BF_Null_ = 3.43, precuneus: BF_Null_ = 3.63, mPFC: BF_Null_ = 4.71). This moderate evidence in support of the null hypothesis highlights the similarity in neural processing of mental states among both autistic and neurotypical individuals.

**Figure 3.**
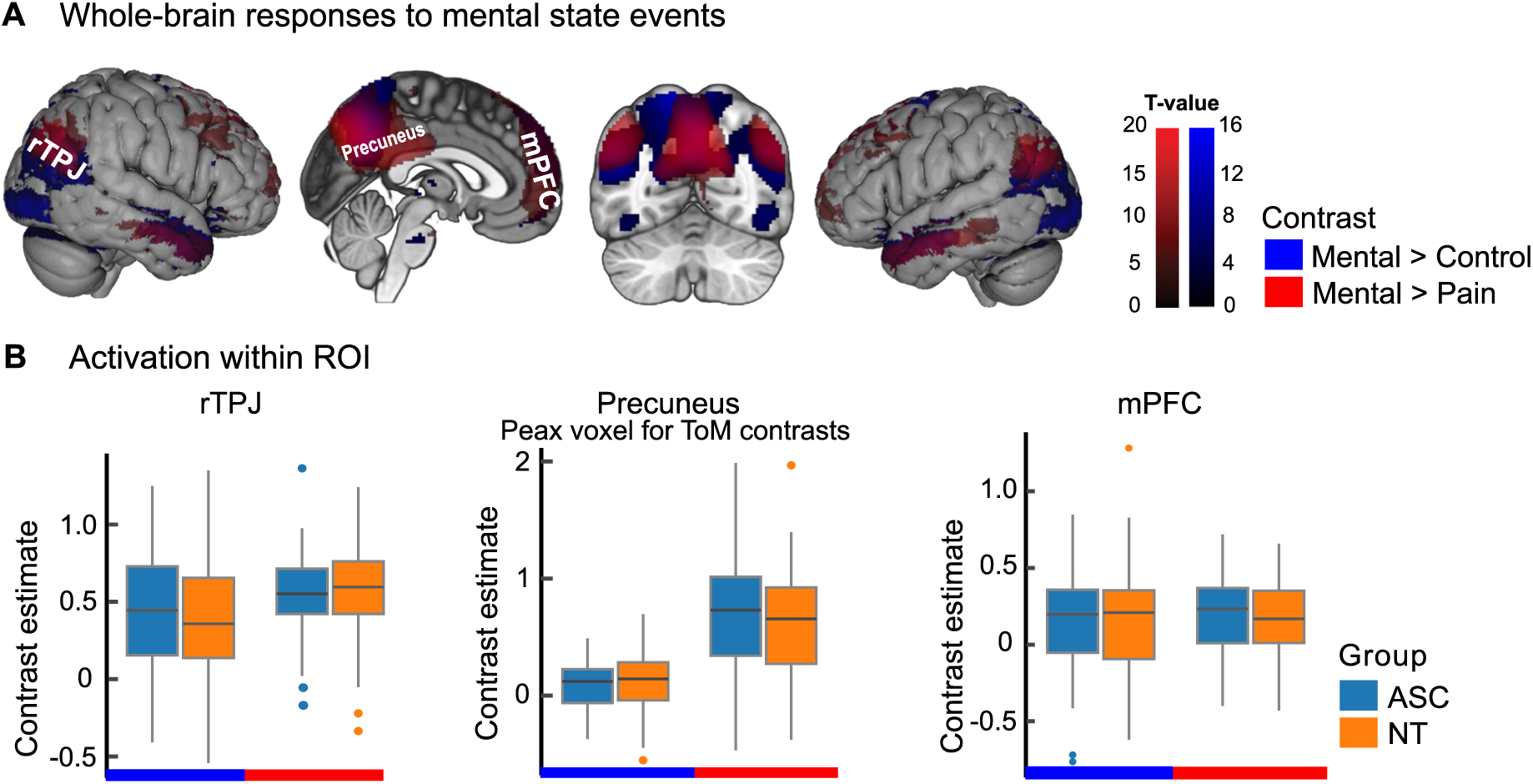
Comparable neural activation patterns during mental state events across groups. (**a**) Brain sections showing whole-brain responses specific to mental state events, as assessed through the Mental > Control and Mental > Pain contrasts. There were no significant differences in activation for either contrast between the autistic and neurotypical groups. (**b**) Box plots depicting the contrast estimates for the Mental > Control (in blue) and Mental > Pain (in red) comparisons across various Regions of Interest (ROI) for both groups. rTPJ and mPFC were selected as ROIs based on ToM literature, while the precuneus was selected because of the peak voxel being located in that region for both ToM contrasts. Across all ROIs and contrasts, no significant differences were observed between the groups. Outliers are represented by dots, while whiskers display a 1.5 inter-quartile range.

### Movie-driven variability in pupil responses

Having observed comparable pupil and brain responses to mental state events, alongside similar verbal descriptions among autistic and neurotypical participants, we aimed to explore differences in narrative processing across the duration of the entire movie. Employing intersubject correlation analysis, we systematically computed correlations across the pupil timeseries using a 30-second sliding window with 100 ms intervals, while also accounting for multiple comparisons across timepoints. This data-driven approach revealed an interval from 40 to 71 seconds, where a significantly stronger intersubject correlation, indicating less variability in pupil responses, was observed in autistic participants compared to neurotypical participants (M_ASC_ = 0.61, M_NT_ = 0.55, *cluster stat* = 777, *p* = .045; Fig. 4). Notably, this interval preceded any mental state events, coinciding instead with the introduction of new characters and a scene of storks flying through clouds. This pattern of reduced variability in autistic participants persisted throughout the movie, although it did not reach statistical significance outside of this interval after adjusting for multiple comparisons. Furthermore, we verified that the observed differences in variability were not due to differences in saccadic eye movements. Analysis indicated that both the frequency and variability of these movements were consistent between autistic and neurotypical participants throughout the movie duration (Fig. S1).

**Figure 4.**
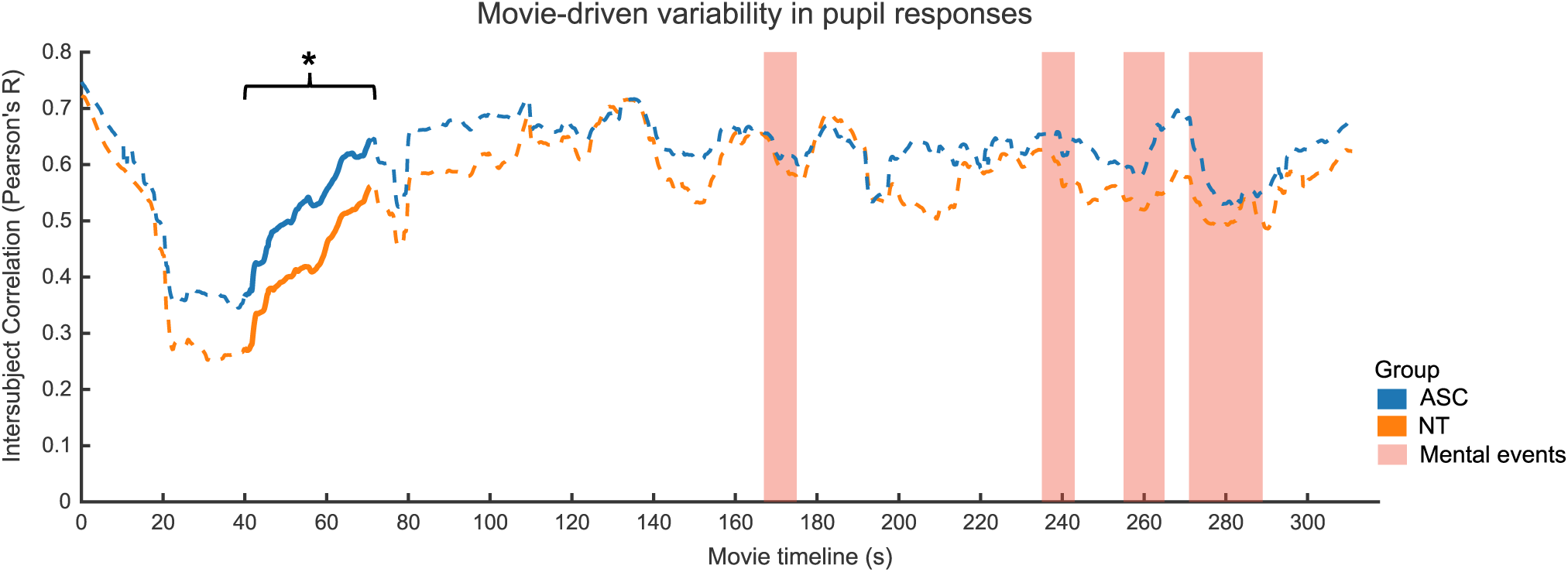
Reduced pupil response variability in autistic participants. Dynamic intersubject correlation analysis indicated comparably high levels of intersubject correlation in both autistic and neurotypical groups at the outset of the movie. However, around 40 seconds in, a significant divergence emerged, with autistic individuals showing stronger intersubject correlations, indicative of less variability, in their pupil responses. This pattern of reduced variability emerged well before the occurrence of mental state events highlighted in red. Solid lines delineate statistically significant intervals, as determined by a cluster-based permutation test.

### Movie-driven variability in brain responses

We proceeded with an intersubject correlation analysis on the fMRI data, employing the same parameters used in the pupillometry analysis. To effectively capture and accommodate potential spatiotemporal variations in brain responses, we enabled the clustering algorithm to dynamically adjust its spatial extent at different timepoints. This analysis identified a significant spatiotemporal cluster where autistic participants showed stronger intersubject correlations, indicative of less variability, in their brain responses compared to neurotypical participants (Fig. 5). This cluster spanned the entire duration of the movie (*cluster stat* = 1332, *p* = .002), and included peaks in several brain regions: the right and left supramarginal gyrus (rSMG: xyz_MNI_ = [52, -34, 32], *t*_max_ = 3.88; lSMG: [-54, -40, 32], *t*_max_ = 4.54), the right inferior temporal gyrus (rITG: [54, -22, -28], *t*_max_ = 5.90), and the left calcarine gyrus (lCG: [6, -102, - 10], *t*_max_ = 4.20). This pattern of reduced variability in brain responses among autistic participants was observed across all peak regions for most of the movie, with transient dips occurring midway and towards its end (Fig. S2). The most pronounced differences were evident in the right and left supramarginal gyrus and the right inferior temporal gyrus, and these differences became apparent following the peak differences in the pupillometry data. Importantly, the cluster emerged well before the mental state events and continued after their conclusion. Furthermore, the cluster had a relatively limited spatial overlap with the ToM network activated during mental state events, comprising less than 20% of the total cluster size (Fig. 6).

**Figure 5.**
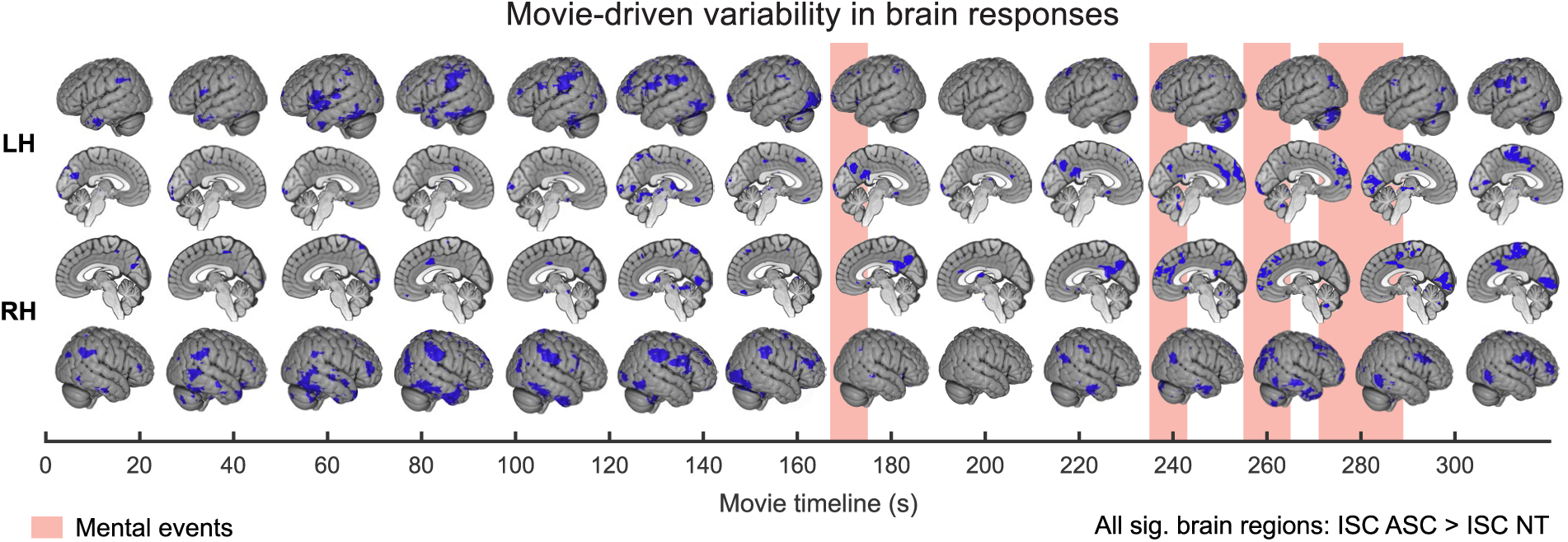
Reduced brain response variability in autistic participants. Dynamic intersubject correlation analysis identified a spatiotemporal brain cluster where autistic participants showed stronger intersubject correlations, indicative of less variability, in brain responses compared to neurotypical participants. This cluster spanned the entire duration of the movie and featured peaks in the right and left supramarginal gyrus, the right inferior temporal gyrus, and the left calcarine gyrus. Echoing the pupillometry data, this pattern of reduced variability in autistic participants emerged well in advance of the mental state events highlighted in red. LH, left hemisphere; RH, right hemisphere.

**Figure 6.**
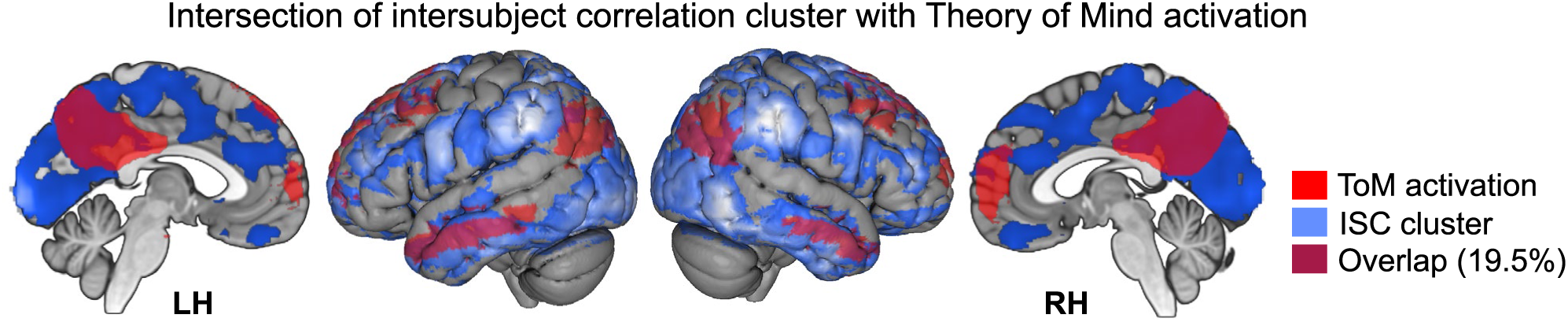
Limited spatial overlap between brain regions showing reduced intersubject variability in autistic participants and Theory of Mind (ToM) activation. The overlap comprised less than 20% of the total cluster size. For clarity, the cluster is visualized using a cumulative t-value threshold of 20 or higher. Lateral brain images provide an overlay with a search distance of 10 mm.

## Discussion

Using fMRI and pupillometry, this study provides functional evidence for the presence of spontaneous mentalizing abilities in high-functioning adults with autism. Compared to neurotypical controls matched for gender, age, verbal and nonverbal IQ, autistic individuals demonstrated similar brain and pupil responses during movie segments known to activate the Theory of Mind network (Jacoby et al., 2016; Richardson et al., 2018). Brain imaging data showed enhanced responses during scenes that encouraged viewers to contemplate the actions and mental states of depicted characters. Conversely, scenes prompting physical state inferences, like characters experiencing physical discomfort, or control scenes without central characters, elicited weaker activations in the ToM network. Participants’ verbal descriptions after the movie reinforced these observations, reflecting a comparable level of engagement with the mental state events depicted in the movie. Collectively, these results extend prior evidence of preserved mentalizing in autism (Dufour et al., 2013; Moessnang et al., 2020), showcasing this capacity in a situation affording but not requiring mentalizing.

While both individuals with and without ASC showed comparable brain and pupil responses during mental state events, intersubject correlation analysis revealed significant differences in the variability of these responses over extended movie intervals. Autistic participants showed reduced variability between individuals across both measures, affecting several brain regions outside the Theory of Mind network. These include the right and left supramarginal gyrus, associated with empathic judgment (Brieber et al., 2007; Ke et al., 2008; Libero et al., 2014), the right inferior temporal gyrus, linked to narrative comprehension (Youssofzadeh et al., 2022), and the left calcarine gyrus, important for visual processing (Woldorff et al., 2002). These differences in response variability were not a simple consequence of discrepancies in bottom-up processing, as both autistic and neurotypical groups demonstrated similar saccadic eye movement patterns. The most pronounced differences and highest variability were detected in early scenes that introduced new characters or depicted a flock of storks in flight. These moments likely encouraged cognitive exploration of the narrative. Research indicates that autistic children may struggle to weave complex causal narratives, possibly reflecting a diminished inclination to place events within a broader narrative context (Losh & Capps, 2003; Tager-Flusberg, 1995). Consequently, the greater variability observed in neurotypical participants could signify a more robust engagement in narrative exploration. The variability in brain regions identified here might be a neural signature of this top-down cognitive processing.

The neuroanatomical bases of the observed changes in response variability among autistic participants are in line with existing research on autism and social interaction. Prior studies have documented changes in the gray matter of the right and left supramarginal gyrus in individuals with autism (Brieber et al., 2007; Ke et al., 2008; Libero et al., 2014), as well as reduced anatomical connectivity in the right inferior temporal cortex (Boets et al., 2018; Koldewyn et al., 2014). The right inferior temporal cortex, known for its sustained activations during narrative comprehension and real-time communication (Stolk et al., 2013; Youssofzadeh et al., 2022), plays a role in the ongoing interpretation and exploration of stimuli. This capability is particularly relevant in everyday social interactions, where interpretations are highly idiosyncratic and fluctuate rapidly as narratives evolve (Goffman, 1974; Johnson et al., 2023; Stolk et al., 2022). Supporting this notion, recent work shows that autistic individuals are less likely to incorporate recent interactional history into their communication, leading to misunderstandings (Wadge et al., 2019). Future research focusing on how stimulus context influences neural variability could shed light on the ways autistic and neurotypical individuals explore and interpret complex narratives.

It is worth noting that our intersubject correlation patterns differ from previous studies demonstrating greater brain response variability in autistic individuals compared to neurotypicals (Byrge et al., 2015; Hahamy et al., 2015; Hasson et al., 2009; Lyons et al., 2020; Nunes et al., 2019; Ou et al., 2022; Pegado et al., 2020; Salmi et al., 2013). Several factors could explain these discrepancies. First, our study used an animated film with fictional characters, as opposed to the more realistic human portrayals in other studies. Though the type of characters may influence brain responses in autism (Atherton & Cross, 2018), greater neural variability has been noted with fictional characters (Lyons et al., 2020). Second, our study implemented an adaptive clustering algorithm to identify spatiotemporal clusters of brain response variability within 30-second intervals. This approach may be more sensitive to brain response variations associated with subtle shifts in interpretation, compared to whole-movie analyses emphasizing consistent patterns over significantly longer durations (10 to 67 minutes). This approach might better capture the theorized heightened reliance on bottom-up sensory stimulation in autism (Pellicano & Burr, 2012), potentially leading to less variability in neural signals related to scene interpretation. More generally, our findings pose challenges for social cognition theories that stress the importance of tight neural synchronization between individuals (Hasson et al., 2012; Mayo et al., 2021). Despite their social challenges, autistic participants in our study showed stronger neural correlations in response to the movie stimuli.

In conclusion, our findings extend prior evidence of preserved mentalizing in autism, showcasing this capacity within a nonverbal, task-free environment. Furthermore, our findings identify a distinct cognitive signature – variability in brain activity and pupil responses to movie narratives. This response variability differentiates neurotypical individuals from those with autism and underscores the importance of narrative exploration in unstructured environments, a situation that is common in everyday social interactions.

## Methods

### Preregistration and Data availability

This study comprised two distinct sets of analyses. The first centered on preregistered comparisons of pupil and brain responses between participant groups, examining specific movie events expected to evoke mental state inferences (Mangnus et al., 2022). The second set of analyses involved computing intersubject correlations to explore potential distinctions in narrative processing throughout the entire movie duration. Unlike the event-related analyses, these exploratory data-driven analyses did not presuppose the relevance of particular segments within the movie. All eyetracking and fMRI data generated during the current study are openly accessible and available for download (Mangnus et al., 2024).

### Participants

Fifty-two high-functioning adults diagnosed with Autism Spectrum Disorder (ASC) and fifty-two matched neurotypical controls (NT) were recruited from Radboud University’s participant database, social media advertisements, campus postings, and local outpatient clinics in Nijmegen and Arnhem (The Netherlands). Inclusion criteria for the ASC group included an official diagnosis from a qualified clinician (American Psychiatric Association, 2013), further validated by the Autism-Spectrum Quotient (AQ-50, Baron-Cohen et al., 2001). Exclusion criteria encompassed the use of psychotropic medication or systemic glucocorticoids, severe cognitive impairment, systemic disease, or neurological treatment history. All participants provided written informed consent following local ethics committee approval (CMO region Arnhem-Nijmegen, The Netherlands, file number 2019-6059) and received study compensation plus travel reimbursements.

The ASC and NT groups did not significantly differ in age (M ± SD 27.7 ± 6.3 vs. 26.0 ± 5.0, *t*(102) = 1.74, *p* = .08, Kullback-Leibler divergence = .05, *F*-test = .29), gender distribution (55.8% vs. 61.5% female, χ^2^(2, 104) = -1.73, *p* = .19), or IQ measured by the Similarities and Vocabulary subscales of the Wechsler Adult Intelligence Scale (WAIS-III, Wechsler, 1997, 126 ± 16 vs. 124 ± 16, *t*(102) = 0.51, *p* = .61, KL = .02, *F*-test = .74) and the Raven’s Progressive Matrices (RPM, Raven, 1989); 103 ± 9 vs. 104 ± 11, *t*(102) = -0.46, *p* = .64, KL = .01, *F*-test = .63). However, autistic participants scored significantly higher on the Autism-Spectrum Quotient (AQ-50, Baron-Cohen et al., 2001; 31 ± 9 vs. 12 ± 6, *t*(102) = 12.2, *p* < .001) compared to neurotypical participants, supporting the distinction between groups. Three participants could not complete the post-movie verbal assessment due to time constraints, and eye-tracking data for two participants was not collected due to technical issues.

### Experimental design

Participants were instructed to view Pixar Animation Studios’ animated movie ‘Partly Cloudy’ while in the scanner (Jacoby et al., 2016). The movie’s duration was 5 minutes and 45 seconds. Following the MRI session, participants completed a questionnaire detailing their interpretation of the movie. They were prompted to describe the storyline in their own words, identify events where the main characters experienced emotions like happiness, sadness, or anger, and discern whether the narrative concluded on a happy or sad note together with their reasoning as to why. The participants were not instructed that they would complete this questionnaire before watching the movie. In this way, the questionnaire was constructed as an unexpected assessment to gauge participants’ grasp of the movie’s plot and their inclination to use language portraying characters’ mental states. Two independent raters classified and tallied words from the verbal descriptions, distinguishing those depicting mental states from other content-related terms based on the list of mental state words in Bang et al. (2013). A subsequent independent *t*-test was employed to compare the usage of words indicative of mental states between the two participant groups.

The movie comprised three distinct event types, coded by the original research team that introduced it as a Theory of Mind localizer task (Jacoby et al., 2016). These events encompassed *Mental*, *Pain*, and *Control* events, each marked along the movie’s timeline in Fig. 1. *Mental* and *Pain* events were expected to elicit inferences about characters’ mental and physical states, while *Control* events involved scenes devoid of foreground characters. For instance, the *Control* condition showcased serene moments like birds in flights or panoramic views of clouds (3 events, totaling 24 seconds). The *Mental* condition provoked viewers to engage with the actions and thoughts of depicted characters, depicting scenarios like characters feeling distressed while observing others enjoy a cheerful interaction or mistakenly perceiving betrayal by a friend (4 events, totaling 44 seconds). Lastly, the *Pain* condition depicted instances where characters experienced physical discomfort, such as being shocked by an eel or bitten by a crocodile (7 events, totaling 26 seconds). These specific events were employed to analyze and contrast both pupillary and neural responses, as detailed in the subsequent sections.

### Pupillometry and MRI data acquisition

Pupil size was continuously tracked using an Eyelink 1000 plus eyetracker at a sampling rate of 1000 Hz. MRI data were acquired using a Siemens 3T MRI-scanner equipped with a 32-channel head coil. Structural images were obtained with a high-resolution T1 MPRAGE sequence with the following parameters: repetition time (TR) = 2200 ms, inversion time (TI) = 1100 ms, echo time (TE) = 2.6 ms, flip angle = 11°, voxel size = 0.8 x 0.8 x 0.8 mm, acceleration factor = 2. Functional images (fMRI) were acquired using a multi-band multi-echo sequence with the following parameters: TR = 1500 ms, number of echos = 3, TE = 13.4/34.8/56.2 ms, flip angle = 75°, voxel size = 2.5 x 2.5 x 2.5 mm, acceleration factor = 2. The ASC and NT groups showed no significant differences in movement, including framewise displacement (FD) across acquired images (mean FD = 0.15 ± 0.05 vs. 0.16 ± 0.09, *t*(102) = - 0.78, *p* = .43; max FD = 0.81 ± 0.79 vs. 1.03 ± 1.31, *t*(102) = -1.09, *p* = .28) and total head motion measured by summing the values of the translation and rotation regressors from the realignment procedure (translation: 109.5 ± 74.6 vs. 114.3 ± 99.5, *t*(102) = 0.28, *p* = .78; rotation: 2.0 ± 0.98 vs. 2.2 ± 1.2, *t*(102) = 0.74, *p* = .46).

### Pupillometry data analysis

Pupillometry data underwent several preprocessing steps, using a combination of established MATLAB routines and custom-developed code. Initially, blinks were identified and removed using a noise-based detection algorithm (Hershman et al., 2018). Following this, a visual inspection was conducted to identify and eliminate squints, characterized by disproportionately small pupil sizes (Mathot, 2018). Additionally, gaze jumps, marked by excessive translational eye movement, were also removed. To identify saccades, we utilized an adaptive velocity threshold algorithm based on the median absolute deviation (Nyström & Holmqvist, 2010). The pupil timeseries data for each participant was then z-scored and adjusted for frame-by-frame global luminance influences. The relationship between luminance and pupil size was modeled using the *lm()* function from the R *stats* package (Bates et al., 2015). In this model, luminance was quantified using RGB values, which were calculated following the relative luminance formula outlined in the Rec. 709 image encoding standard (ITU, 2002). We employed polynomial functions up to the 5th order, which demonstrated improved fitting after evaluation with five randomly selected participants. The residuals obtained from this modeling were used for subsequent pupillometry data analysis. Importantly, it was confirmed that this luminance correction did not alter the results of our group analyses relating to pupil sizes and their variability.

To examine variations in pupil size, which are considered an index of cognitive processing effort (Beatty, 1982), we employed a 3x2 mixed-design ANOVA using mean pupil size as the dependent variable. The within-subject factor comprised event conditions (*Mental*, *Pain*, *Control*), while participant group status (ASC, NT) served as the between-subject factor. We further employed Tukey’s Honest Significant Difference tests to examine ToM-related contrasts (*Mental* > *Pain*, *Mental* > *Control*). To examine differences in intersubject variability driven by the movie’s narrative, we conducted an intersubject correlation analysis on the pupil timeseries. Employing a leave-one-out approach, we calculated each participant’s timeseries correlation with the average of all other group members’ timeseries (Nastase et al., 2019). Using a 30-second sliding window at 100 ms intervals, this method generated a correlation timeseries across the entire movie duration for each participant. Subsequently, all Fisher *z*-transformed correlation timeseries were subjected to a nonparametric cluster-based permutation test (Maris & Oostenveld, 2007) evaluating differences in pupil response variability between the participants groups, while correcting for multiple comparisons across timepoints (two-sided independent samples *t*-test, 10,000 permutations, Monte-Carlo *p* < .05).

### fMRI data analysis

Functional images underwent a series of preprocessing steps implemented in SPM12 (https://www.fil.ion.ucl.ac.uk/spm/). Initially, the echoes were consolidated into a single volume through echo-weighted combination. Subsequent steps involved realignment using 2nd degree B-spline interpolation and six rigid-body transformation parameters to align the images with the first image. To minimize spatial distortions and signal dropout, unwarping with an acquired fieldmap was applied. Anatomical images were coregistered with the mean functional images. Next, segmentation using SPM’s tissue probability maps categorized anatomical tissue into gray matter, white matter, and cerebral spinal fluid, facilitating normalization to MNI space for the functional images. Following normalization, a spatial smoothing kernel of 8 mm full-width at half-maximum was applied. First-level regressors accounted for *Mental*, *Pain*, and *Control* events of the movie, alongside regressors for the end credits, head movement (including squared and cubic terms, and first and second derivatives), and signal intensities from different tissue compartments. Finally, a 0.6 threshold masking was applied for optimal brain inclusion.

Group differences in brain responses to the different events were examined via two 2-by-2 mixed-design ANOVAs, focusing on Theory of Mind (ToM) processing. These analyses used group status (ASC, NT) as the between-subject factor and considered either *Mental* and *Pain* events or *Mental* and *Control* events as within-subject factors. The former ToM contrast aimed to replicate the stronger activation of the ToM network, as found in Jacoby et al. (2016). The latter contrast aimed to explore the generalization of these findings to scenes without foreground characters, i.e. control events. We report whole-brain cluster-level corrected findings (*p_FWE_* < .05), with anatomical locations in gray matter determined using the probabilistic atlas from the SPM Anatomy Toolbox (Eickhoff et al., 2005). Additionally, we conducted a region of interest (ROI) analysis centered on three peak brain regions identified by the two ToM contrasts. To rigorously test comparable neural activation in the ToM network between ASC and NT participants, we employed Bayesian analysis (JASP Team, 2022). This approach computes Bayes Factors (BFs) to assess the evidence favoring the null hypothesis over models that differentiate between autistic and neurotypical participants.

To examine differences in intersubject variability at the neural level, we conducted an intersubject correlation analysis on brain voxel timeseries accounted for head movement and tissue compartment signal intensities. Employing a leave-one-out approach mirroring the pupillometry analysis, we explored dynamic spatial variations in brain responses. Our adaptive clustering algorithm sought spatiotemporal clusters indicating distinct brain response variability between ASC and NT participants. For computational efficiency, we downsampled the fMRI data spatially and temporally by a factor of 3. Lastly, we subjected all spatiotemporal patterns to a nonparametric cluster-based permutation test (Maris & Oostenveld, 2007), correcting for multiple comparisons across voxels and timepoints (two-sided independent samples *t*-test, 10,000 permutations, Monte-Carlo *p* < .05). To assess the spatial distribution of the identified spatiotemporal brain cluster, we summed its t-values reflecting variability differences across the entire movie, excluding summed t-values below 20. To quantify the degree of overlap between this brain correlation cluster and the ToM network, activated during mental state events, we calculated the proportion of their spatial overlap in relation to the total voxel size of the brain correlation cluster.

## Supplementary Information

**Table S1.** Results of the within-subject fMRI analyses related to the main effects of event type (Mental, Pain, Control).

| *Contrast* | *Anatomical location of peak voxel* | *Cluster size* | *MNI coordinates* |  |  | *T-value*<br>(df = 1, 204) |
| --- | --- | --- | --- | --- | --- | --- |
|  |  |  | *x* | *y* | *z* |  |
| Mental > Pain | Right precuneus | 19445 | 6 | -64 | 40 | 20.64 |
|  | Right superior frontal gyrus | 16376 | 28 | 26 | 54 | 12.76 |
|  | Left middle temporal gyrus | 2560 | -52 | 2 | -26 | 11.51 |
|  | Right middle temporal gyrus | 1782 | 60 | -12 | -20 | 10.14 |
|  | Right parahippocampal gyrus | 311 | 22 | -40 | -10 | 5.40 |
| Mental > Control | Left precuneus | 31320 | -2 | -54 | 44 | 15.25 |
|  | Left middle temporal gyrus | 2525 | -56 | -8 | -20 | 10.58 |
|  | Left superior medial gyrus | 8939 | -6 | 52 | 38 | 7.33 |

**Figure S1.**
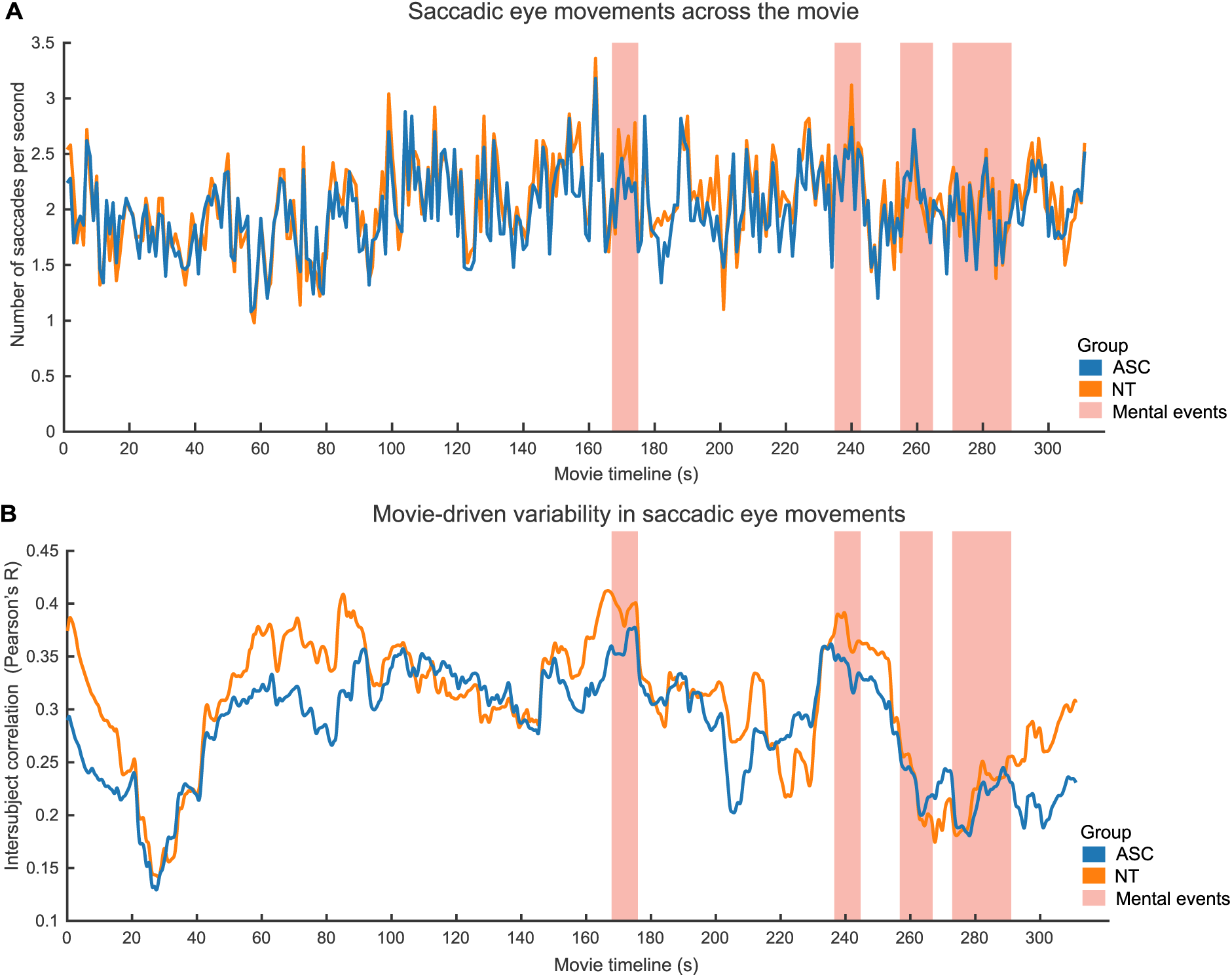
Frequency and variability of saccadic eye movements. (**a**) Mean number of saccades observed throughout the movie in both autistic and neurotypical individuals. No significant differences in the frequency of saccades were detected between the two groups. (**b**) Mean intersubject correlation coefficients representing the variability in the number of saccades across the movie among autistic and neurotypical individuals. No significant differences in saccade variability were observed between the two groups.

**Figure S2.**
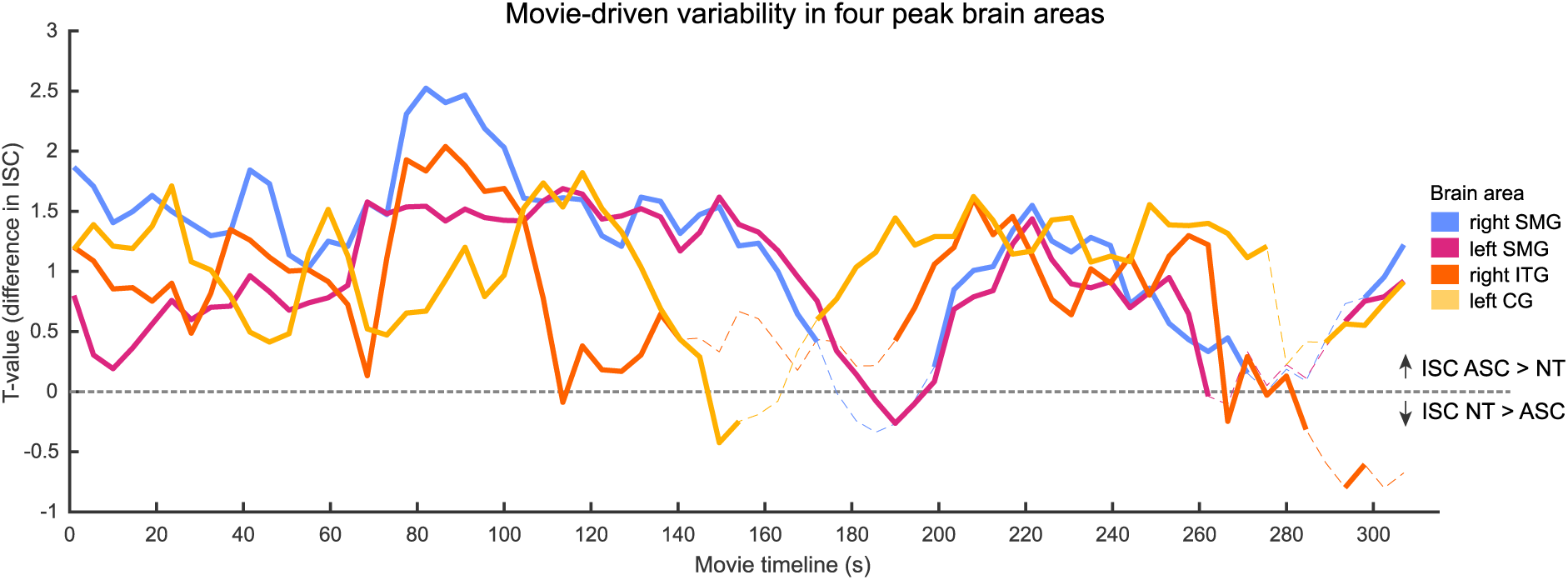
Brain response variability across four key regions, including the right and left supramarginal gyrus (rSMG; lSMG), the right inferior temporal gyrus (rITG), and the left calcarine gyrus (lCG). T-values were extracted from a 30 mm diameter sphere centered on the peak voxel. Variability in all four regions was statistically significant throughout most of the movie duration.

